# Bone Structural, Biomechanical and Histomorphometric Characteristics of the Postcranial Skeleton of the Marsh Rice Rat (*Oryzomys palustris*)

**DOI:** 10.1101/2021.09.02.458784

**Authors:** E. J. Castillo, S. Croft, J.M Jiron, J.I. Aguirre

**Author notes:** Please address correspondence to: J. Ignacio Aguirre, D.V.M., Ph.D., DACLAM, Department of Physiological Sciences, Box 100144, JHMHC, University of Florida, Gainesville, FL 32610.

## Abstract

**INTRODUCTION:** The rice rat (*Oryzomys palustris*) is a non-conventional laboratory rodent species used to model some human bone disorders. However, no studies have been conducted to characterize the postcranial skeleton. Therefore, we aimed to investigate age- and gender-related features of the appendicular skeleton of this species.

**METHODS:** We used femurs and tibiae from 94 rats of both genders aged 4-28 wks. Bone mineral content (BMC), bone mineral density (BMD), and biomechanical properties were determined in femurs. In addition, bone histomorphometry of tibiae was conducted to assess bone cells activities and bone turnover over time.

**RESULTS:** Bodyweight, bone length, total metaphysis BMC/BMD, cortical BMC/BMD, cortical thickness, and cortical area progressively augmented with age. Whereas the increase in these parameters plateaued at age 16-22 wks in female rats, they continued to rise to age 28 wks in male rats. Furthermore, bone strength parameters increased with age, with few differences between genders. We also observed a rapid decrease in longitudinal growth between ages 4-16 wks. Whereas young rats had a greater bone formation rate and bone turnover, older rice rats had greater bone volume and trabecular thickness, with no differences between genders.

**CONCLUSIONS:** 1) Sexual dimorphism in the rice rat becomes grossly evident at age 16 wks; 2) the age-related increases in bone mass, structural cortical parameters, and in some biomechanical property parameters plateau at an older age in male than in female rats; and 3) bone growth and remodeling significantly decreased with age indistinctive of the gender.

## Introduction

*Cricetidae* is the most prominent family of rodents, and the second-largest family of mammals, with 698 species that include the rice rat, hamster, vole, lemming, and new world mice and rats (Eisenberg, 1984). Rodents in this family are very diverse and range in size from tiny mice to muskrats. *Cricetidae* are found in various habitats, including swamps, wetlands, tundra, deciduous forests, coniferous forests, rainforests, deserts, mountains, and urban areas from North America, South America, Europe, and Asia. The marsh rice rat (*Oryzomys palustris*) is taxonomically classified within the Order: Rodentia; Suborder: Myomorpha; Family: Cricetidae; Subfamily: Sigmodontinae; Tribe: *Oryzomyini;* Genus: *Oryzomys;* Species: *palustris* (Weksler and Percequillo, 2011; Weksler, 2014; Weksler et al., 2014). Marsh rice rats can be found inhabiting the salt marshes and wetlands of North and South America (Hamilton, 1946) due to their well-adapted conditions for swimming (Esher et al., 1978). Marsh rice rats are medium-sized rodents with a total length of 226-305 mm and substantial sexual dimorphism (Goldman, 1918; Hamilton, 1946; Hall and Kelson, 1959; Hamilton and Whitaker, 1979; Wolfe, 1982).

Some species within *Cricetidae* have been adapted to live in laboratory-reared environments, including the hamster, voles, deer mouse, and rice rat. We established one of the only two rice rat colonies in the US at the University of Florida (UF) in 2013 (Aguirre et al., 2015).

The rice rat is a USDA-covered species housed and maintained according to the Animal Welfare Act (AWA) regulations and PHS Policy, ensuring societal aspects of individual species and enabling scientifically sound research (**Figure 1**). The rice rat was initially used in research after transitioning from the wild to captivity in the 1950s (Gupta and Shaw, 1956b, 1956a). Throughout decades, they have been used to study physiologic processes and various pathologic disorders. Indeed, the rice rat has been used to study photoperiod and the pineal gland’s role in reproduction and reproductive development (Edmonds and Stetson, 1993, 1994; Edmonds et al., 1995; Edmonds and Stetson, 2001; Edmonds et al., 2005). They have also been utilized for studying the airway epithelium and disclose the distinctive transport properties that these species possess in their airways, which may explain their ability to survive in marsh salt environments (Kuan et al., 2019). The rice rat also appears to be a unique species for studying retinal pigmented epithelium degeneration (Casavant et al., 2000; Grahn et al., 2005).

**Figure 1.**
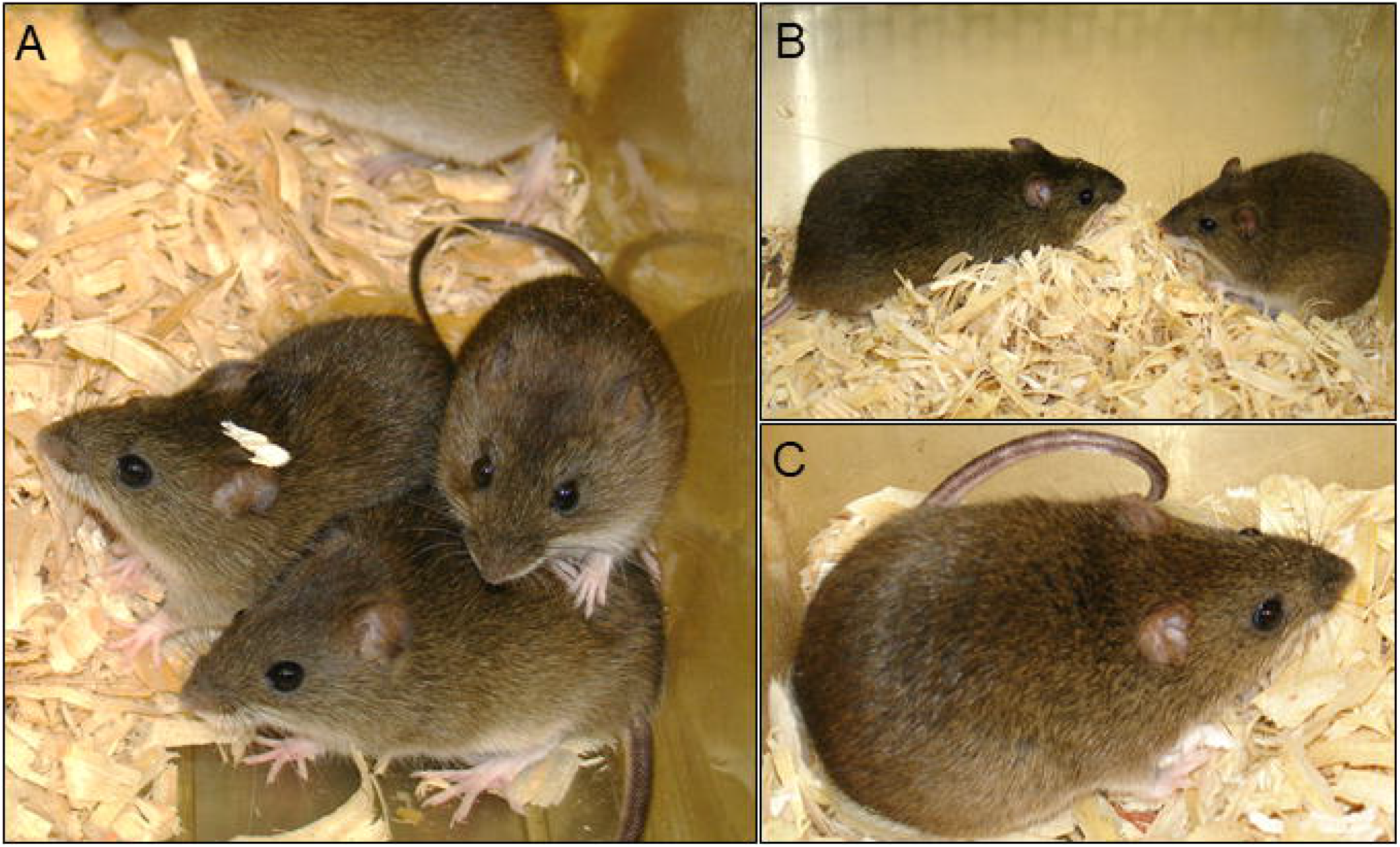
Rice rats under laboratory animal conditions. **A**. Rice rats are housed 2-5 per group of the same sex and placed in static filter top cages with pine shavings as bedding. **B**. Male and female rice rats are paired together during the breeding period. Note the significant sexual dimorphism between a male (left) and a female rice rat (right). **C**. Pregnant female in a breeding cage.

Furthermore, rice rats have been extensively used in periodontal disease research since it is a species extraordinarily susceptible to the initiation and progression of periodontitis, with accelerated alveolar bone loss, without requiring any intraoral mechanical manipulation (Gupta and Shaw, 1956b, 1956a; Ryder, 1980; Gotcher and Jee, 1981; Aguirre et al., 2012a). In addition, they have been used to establish a reliable model of medication-related osteonecrosis of the jaw (MRONJ) (Aguirre et al., 2012b; Messer et al., 2018; Messer et al., 2019; Castillo et al., 2021). MRONJ is a potentially severe, debilitating oral condition characterized by exposed, necrotic bone in the maxillofacial region of patients with cancer and patients with osteoporosis who have been taking antiresorptive and/or antiangiogenic medication to reduce bone metastases and skeletal fractures, respectively (Ruggiero et al., 2014; Khan et al., 2015).

While studying periodontitis and MRONJ in this species, various features of craniofacial bone growth, modeling and remodeling have been described (Aguirre et al., 2012b; Messer et al., 2017; Messer et al., 2018; Messer et al., 2019; Messer et al., 2020; Castillo et al., 2021). Though, little attention has been given to the postcranial skeleton, particularly the appendicular bones. Preliminary studies from our group have noticed a remarkable response not only by craniofacial bones of this species but also by appendicular bones to antiresorptive and bone anabolic agents (Aguirre et al., 2012b; Messer et al., 2018; Messer et al., 2019). However, no formal studies have been conducted to investigate the postcranial skeleton’s baseline physiologic and phenotypic characteristics in this species. Thus, this study aimed to investigate age-related and gender-specific features of the appendicular skeleton, evaluating quantitative skeletal endpoints using bone histomorphometry and peripheral quantitative computed tomography (pQCT), and biomechanical property strength testing.

## Materials and Methods

### Experimental rice rats and animal care

Femurs and tibiae were used from 94 rice rats (40 males and 54 females) age 4 to 28 wks. Rice rats were generated in-house utilizing a monogamous continuous breeding system (Aguirre et al., 2015) (**Figure 1**). Animal care and experimental procedures were conducted following federal policies and guidelines of the University of Florida Institutional Animal Care and Use Committee (IACUC). Adequate measures were taken to minimize pain and discomfort in the rats. The Animal Care Services resource at the University of Florida is an AAALAC-accredited animal care and use program.

### Euthanasia and tissue collection

Rice rats were injected SC with declomycin and calcein (Sigma Chemical Co., St Louis, MO) at a dose of 15 mg/kg on the 7^th^ and 2^nd^ days before sacrifice, respectively, to label sites of bone formation for the bone histomorphometry analysis (Aguirre et al., 2001; Aguirre et al., 2007; Aguirre et al., 2012b). Rice rats were euthanized by CO_2_ inhalation followed by cervical dislocation. Femurs and tibiae were stripped of musculature, leaving the periosteum intact. The length of these bones was measured using a digital caliper. Immediately after this procedure, tibiae were fixed in 10% phosphate-buffered formalin for 48 hr, placed in 70% ethanol, and processed for cancellous bone histomorphometry. The femurs were used to determine bone mineral content (BMC) and bone mineral density (BMD) by peripheral quantitative computed tomography (pQCT) (see below), then wrapped in saline-soaked gauze and stored at -20°C until subsequent bone biomechanical testing (see below).

### Bodyweight and measurements of bone length

Bodyweight measurements in rice rats of both genders were taken every two weeks from age 4 wks to age 28 wks. The length of femurs and tibiae were taken blind using Fowler calipers (Fowler Sylvac, ultra-cal mark III). The femur length was taken from the neck of the proximal epiphysis, between the head and the greater trochanter, to the surface of the distal epiphyseal condyles. The tibial length was taken from the intercondylar eminence of the proximal epiphysis to the medial malleolus of the distal epiphysis.

### Peripheral quantitative computed tomography (pQCT) analysis

The pQCT analysis was conducted in femurs of rice rats of both genders at age 4 wks, 10 wks, 16 wks, 22 wks, and 28 wks. Femurs were thawed to room temperature and remained wrapped in saline-soaked gauze except during scanning using a Stratec XCT Research M instrument (Norland Medical Systems; Fort Atkinson, WI), using the company software version 5.40. Scans were taken at a distance of 5 mm proximal to the distal end of the femur (distal epiphysis) and at midshaft. The former site is located at the level of the secondary spongiosa of the distal femoral metaphysis. The midshaft site is located at 50% of the average height of the femur corresponding to the cortical bone of the diaphysis. Volumetric content, bone density, and bone area were determined for total bone (trabecular and cortical bone) at the distal metaphysis and midshaft as previously described (Ke et al., 2001).

### Bone biomechanical properties

After pQCT analysis, femurs were subjected to 3 point-bending strength tests using a material testing system (Bose ElectroForce 3220, Eden Prairie, MN, USA) with a constant span of 9.6 mm as previously described (Ko et al., 2018). Briefly, femurs were placed horizontally, centered on the supports, and directly vertical to the midshaft. Femurs were tested at a constant displacement of 0.05 mm/s until failure. The applied force was measured with a 225 N load cell. Test data were sampled at 2500 Hz and analyzed for stiffness (N/mm), ultimate load (N), work to fracture (Nmm), elastic modulus (Mpa), and ultimate stress (Mpa) using appropriate calculations derived from the load-displacement curve. In addition, bivariate Pearson’s correlation coefficient was calculated between pQCT and biomechanical parameters to determine any relationships between the continuous variables. The interpretation of Pearson’s correlation coefficient, R, was based on previous studies (Mukaka, 2012). Thus, values between 0.90-1; 0.70-0.90; 0.50-0.70; 0.30-0.50; and 0.0-0.30 were considered as very high, high, moderate, low, and negligible correlations, respectively.

### Bone histomorphometry

Tibial bone growth and cancellous bone histomorphometry parameters were assessed at the proximal tibial metaphysis using standard histomorphometry techniques with the Osteomeasure System (Osteometrics, Decatur, GA, USA) (Aguirre et al., 2001; Iwaniec et al., 2007). In brief, tibiae were cut cross-sectionally at the midshaft with a Dremel Moto-Tool (Dremel, Racine, WI), dehydrated in ethanol, and embedded undecalcified in methyl methacrylate (Baron et al., 1983). Proximal tibiae were sectioned longitudinally at 4- and 8-μm thicknesses with a Leica/Jung 2265 microtome. The 4-μm bone sections were stained with the von Kossa method and counterstained with tetrachrome (Polysciences Inc., Warrington, PA, USA). The 8-μm sections remained unstained to measure fluorochrome-based indices of bone formation.

Tibial bone growth was assessed by determining the longitudinal growth rate (LGR) in 8 µm unstained sections at a magnification of 100x. The LGR was calculated by measuring the distance between the growth plate-metaphyseal junction and the fluorescent calcein bands that parallels the growth plate at the primary spongiosa at six equally spaced sites per section. This distance was then divided by the time interval between the administration of calcein and the day of euthanasia (2 days) (Li et al., 1996).

For cancellous bone histomorphometry, measurements were performed within a region of interest (ROI) in the secondary *spongiosa*. The ROI was defined between 400 µm (excluding the primary spongiosa) to 1500 µm from the growth plate, staying away from the endocortical surfaces by ∼250 µm. Structural parameters were measured in von Kossa stained sections at a magnification of x40. Structural parameters included cancellous bone volume (BV/TV), trabecular thickness (Tb. Th), and trabecular number (Tb.N). Static histomorphometric parameters were also measured in the von Kossa stained sections but at a magnification of x200. Static parameters included osteoblast (Ob.S/BS) and osteoclast (Oc.S/BS) surfaces, as percentages of the total cancellous perimeter and number of osteoblasts (N.Ob/B.Pm) and osteoclasts (N.Oc/B.Pm). Dynamic histomorphometry was conducted to evaluate different parameters using fluorochrome-based indices of cancellous bone formation in 8 µm thick, unstained sections at x200 under ultraviolet illumination. Single-labeled surfaces (sL/BS) and double-labeled surfaces (dL/BS) are indices of bone formation, where bone surfaces displayed one or two fluorochrome labels. Mineralizing surface (MS/BS), an index of active bone formation, was calculated as the percentage of the cancellous bone surface that displayed double-labeled surfaces plus one-half of the single-labeled surface. Mineral apposition rate (MAR), an index of osteoblast activity, was calculated by dividing the interlabel distance by the time interval between administration of fluorochrome labels. Bone formation rate (BFR/BS) was calculated by multiplying MS/BS by MAR (Frost, 1983). The terminology used was based on recommendations of the Histomorphometry Nomenclature Committee of the American Society of Bone and Mineral Research (Parfitt et al., 1987; Dempster et al., 2013).

### Statistical analysis

Data are expressed as mean ± SD for each group and evaluated with ANOVA followed by the Holm-Sidak test for multiple comparisons. When ANOVA assumptions regarding normality of data were not met, the non-parametric Kruskal-Wallis test was used. Regardless of the test employed, *P* values less than 0.05 were considered to be statistically significant. To establish the strength and direction of linear relationships between the pQCT and biomechanical parameters data, we applied Bivariate Pearson correlation using the correlation function in excel with two-tailed distribution to determine *P* values.

## RESULTS

### Bodyweight and measurements of bone length

Rice rats progressively grew from weaning (age 4 wks) to adulthood (age 14 wks), as demonstrated by the sustained increase in body weight gain in both males and females (**Figure 2 A**). The gain in weight was indistinguishable between genders up to age 14 wks, with no significant differences between males and females during this period. Whereas male rice rats’ body weight continued to increase from age 14 wks to age 28 wks, bodyweight plateaued at age 16 wks in female rice rats. As a consequence of these changes, male rice rats weighed more than female rats from age 16 wks to age 28 wks (p < 0.005) (**Figure 2 A**).

**Figure 2.**
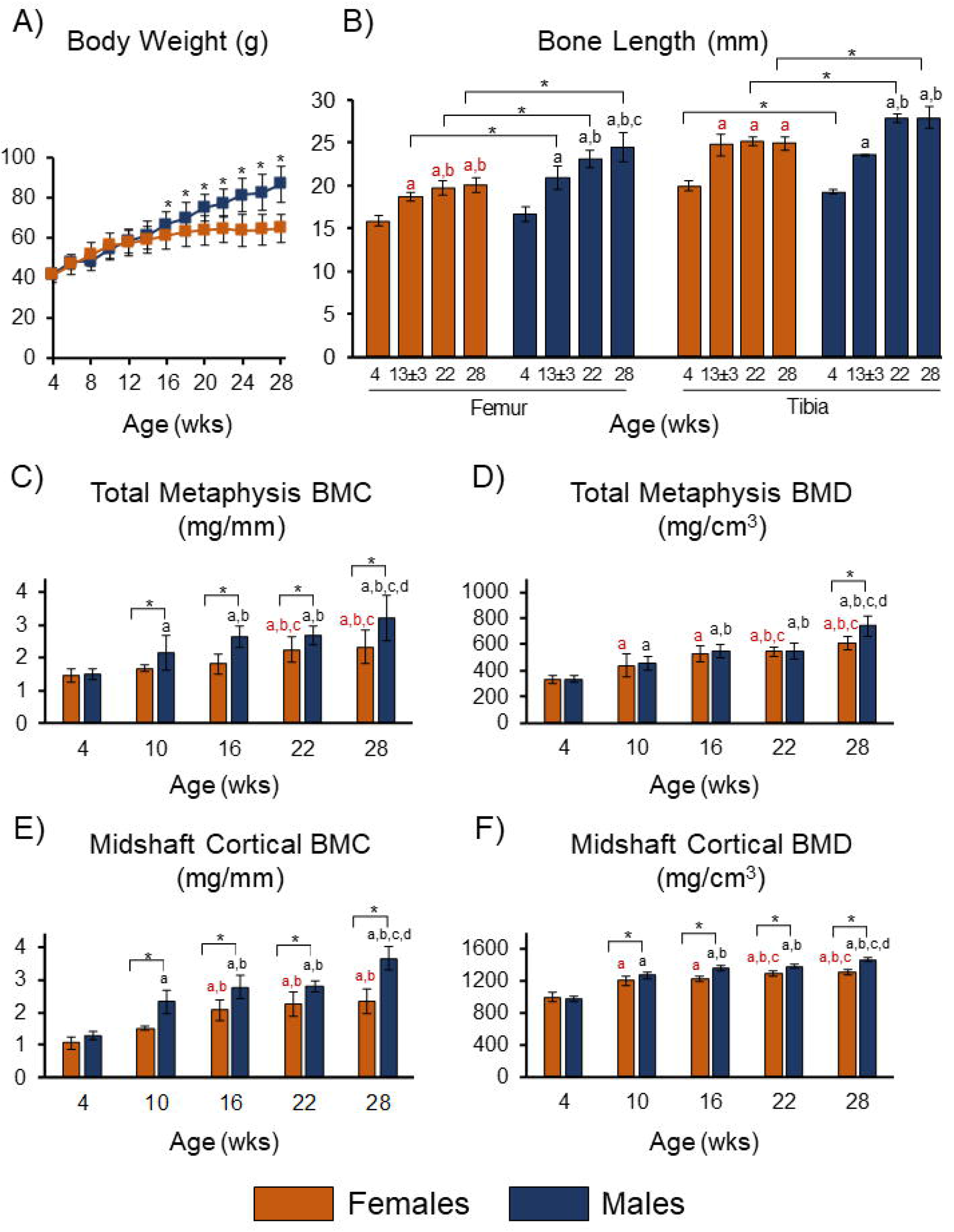
Bodyweight, bone length, and femoral pQCT parameters. **A**. Bodyweight in rice rats from age 4 to 28 wks. **B**. Femoral and tibial bone length in rice rats from age 4 wks to age 28 wks. ^a-c^ denotes significant differences within each gender (p<0.05). **C**. Total distal femur metaphysis bone mineral content (BMC) in rice rats of both genders from age 4 to 28 wks. **D**. Total bone mineral density (BMD) of the distal femur metaphysis in rice rats of both genders from age 4 to 28 wks. **E**. Femur midshaft cortical BMC in rice rats of both genders from age 4 to 28 wks. **F**. Femur midshaft cortical BMD in rice rats of both genders from age 4 to 28 wks. ^a-d^ denotes significant differences within each gender (p<0.05). *indicates significant differences between genders (p<0.001). Bars represent means ± SD.

Consistent with the bodyweight data, femurs and tibiae increased in length from age 4 wks to 13 ± 3 wks, with comparable values for males and females (**Figure 2 B**). Whereas the length of femurs and tibiae progressively increased in male rice rats after age 13± 3 wks, the size tended to plateau at age 13 ± 3 wks in female rats. The mean femur length was 11%, 15%, and 18% greater in male than in female rats at age 13 ± 3 wks (p<0.001), 22 wks (p<0.001), and 28 wks (p<0.001), respectively. The mean tibial length was 4% shorter at age 4 wks (p=0.046), but 10% and 11% greater in male rats than female rats at age 22 wks (p<0.001) and age 28 wks (p<0.001), respectively (**Figure 2 B**).

### pQCT analysis

In consonance with the steady increase in body weight and length of the appendicular skeleton with age, we observed a progressive augmentation in total metaphysis BMC and BMD at the distal femur in rice rats of both genders (**Figures 2 C&D**). Total metaphysis BMC and BMD were greater in older females (age 22-28 wks) than in younger female rats age 4 wks, 10 wks, and 16 wks, respectively (P<0.05). However, no differences in these parameters were found between females age 22 and 28 wks. In male rice rats, these parameters were significantly greater at age 28 wks than all the other age groups (p<0.05). Furthermore, two-way ANOVA showed a greater total metaphysis BMC from age 10 wks to age 28 wks (p<0.001), but only at age 28 wks (p=0.037) in total metaphysis BMD in male rats compared to female rats.

Cortical BMC and BMD tended to increase with age in rice rats of both genders (**Figures 2 E&F**). Specifically, in female rats, cortical BMC progressively increased from age 4 wks to age 16 wks. However, no differences were found between ages 16-28 wks. Similarly, cortical BMD increased from age 4 to 22 wks, but no differences were found between ages 22 wks and 28 wks. In male rice rats, both cortical BMC and BMD progressively increased from age 4 wks to age 28 wks. Two-way ANOVA showed that male rice rats had significantly greater cortical BMC and BMD from age 10 wks to age 28 wks (p<0.001) than age-matched female rats.

The midshaft cortical bone area and cortical bone thickness also tended to increase with age in both genders (**Table 1**). In female rice rats, the midshaft cortical area was greater at age ≥16 wks than in younger rats age ≤10 wks, and cortical thickness increased from age 4 to 22 wks (p<0.05). In male rice rats, both midshaft cortical bone area and cortical thickness tended to increase with age more consistently (p<0.05) (**Table 1**). Whereas no differences were found in the moment of inertia with age in female rats, in male rats, this parameter was significantly greater at age 28 wks compared to all the other age groups (p<0.05) (**Table 1**).

**Table 1.** Structural cortical bone data of rice rat femurs distributed by gender and age.

| **Females** |  |  |  |  |  | **Males** |  |  |  |  |
| --- | --- | --- | --- | --- | --- | --- | --- | --- | --- | --- |
| **Age (wk-old)** | **4 (a)** | **10 (b)** | **16 (c)** | **22 (d)** | **28 (e)** | **4 (A)** | **10 (B)** | **16 (C)** | **22 (D)** | **28 (E)** |
| **Midshaft cortical area (mm<sup>2</sup>)** | 1.07<br>±0.14 | 1.25<br>±0.05 | 1.63<sup>ab</sup><br>±0.15 | 1.73<sup>ab</sup><br>±0.21 | 1.77<sup>ab</sup><br>±0.24 | 1.29*<br>±0.21 | 1.83<sup>A*</sup><br>±0.35 | 2.03<sup>A*</sup><br>±0.33 | 2.08<sup>AB*</sup><br>±0.23 | 2.49<sup>ABCD*</sup><br>±0.22 |
| **Midshaft cortical thickness (mm)** | 0.24<br>±0.02 | 0.33<sup>a</sup><br>±0.07 | 0.42<sup>ab</sup><br>±0.05 | 0.47<sup>abc</sup><br>±0.05 | 0.50<sup>abc</sup><br>±0.05 | 0.29*<br>±0.02 | 0.39<sup>A*</sup><br>±0.03 | 0.45<sup>AB</sup><br>±0.04 | 0.55<sup>ABC*</sup><br>±0.02 | 0.64<sup>ABCD*</sup><br>±0.03 |
| **Moment of inertia (mm<sup>4</sup>)** | 0.30<br>±0.10 | 0.33<br>±0.08 | 0.43<br>±0.07 | 0.37<br>±0.1 | 0.35<br>±0.08 | 0.33<br>±0.07 | 0.53<sup>A*</sup><br>±0.12 | 0.59<sup>A*</sup><br>±0.11 | 0.57<sup>A*</sup><br>±0.12 | 0.73<sup>ABCD*</sup><br>±0.14 |

Femoral midshaft morphology was determined by pQCT. Data are presented as means ± standard deviations. \*Significantly different from the respective female age-matched group (P<0.05). Superscript letters a-e indicate significant differences in female rice rats from the respectively labeled age group (P<0.05). Superscript letters A-E indicate significant differences in male rice rats from the respectively labeled age group (P<0.05).

Two-way ANOVA estimated gender effects, with male rats having a greater midshaft cortical bone area and cortical thickness at almost every age-time point (p<0.05) (**Table 1**). Furthermore, femurs of male rats had a greater moment of inertia (p<0.05) than female rats at age ≥10 wks.

### Bone biomechanical properties

Three-point bending of the femoral midshaft showed a significant age-related increase in stiffness (p<0.001) and ultimate load (P<0.001) in rice rats, regardless of gender (**Table 2**). Work to fracture in female rice rats was greater at age 28 wks than at age ≤10 wks (p<0.001). In male rice rats, work to fracture was greater at age 28 wks than at age wks (p=0.002) and 22 wks (p=0.005).

**Table 2.**
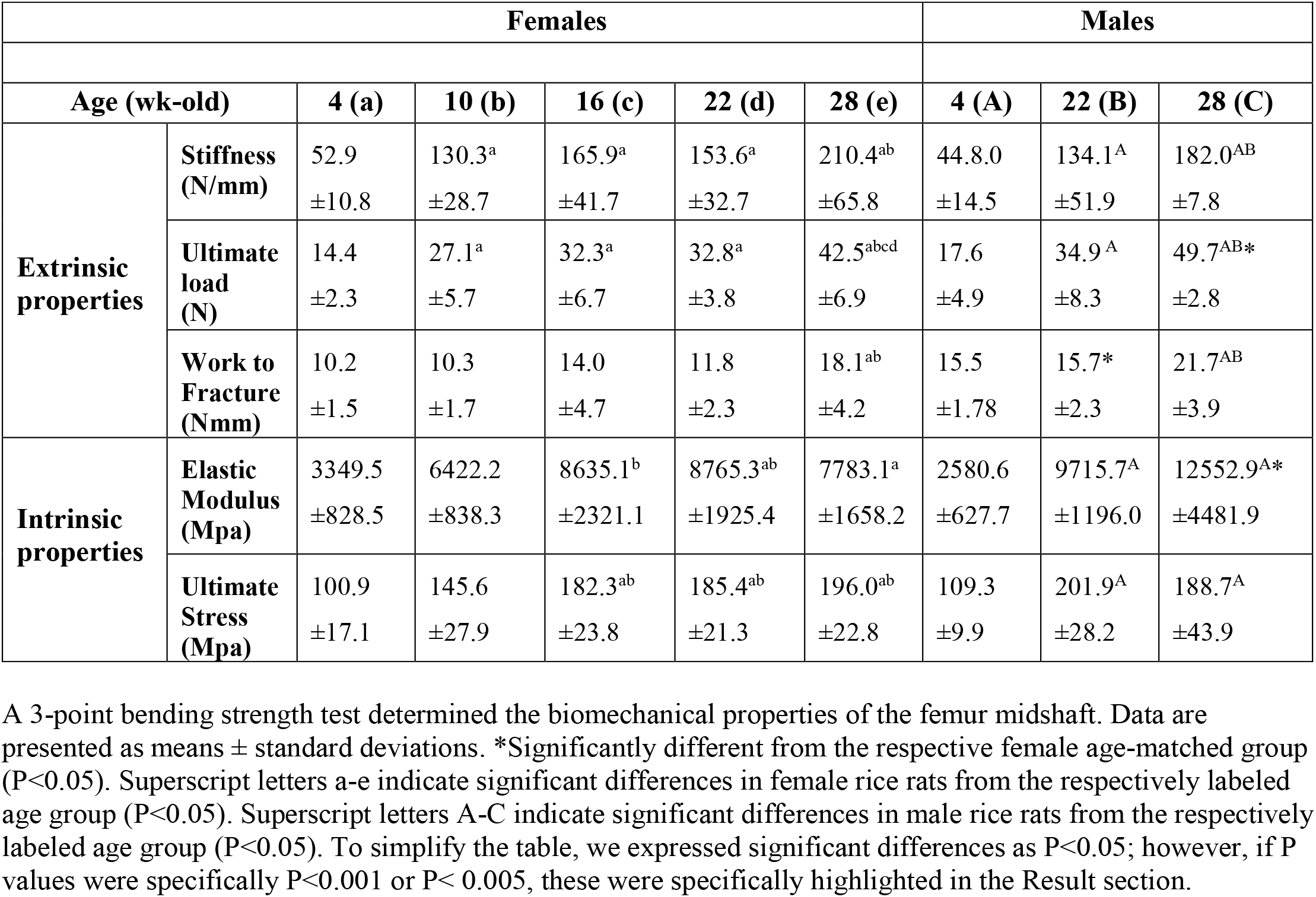
Biomechanical properties data of rice rat femurs distributed by gender and age.

Furthermore, we found an age-related increase in a few of the intrinsic properties of the femoral midshaft. Indeed, the elastic modulus was more significant in older rice rats (age 22-28 wks) compared to age 4 wks rats, regardless of gender (p<0.001) (**Table 2**). In addition, ultimate stress was greater in female rats age ≥16 weeks than female rats age ≤10 wks (p<0.05). However, no differences among age groups were found beyond age 16 wks. Similarly, ultimate stress was significantly greater in older male rice rats (age 22-28 wks) compared to rats age 4 wks.

The two-way ANOVA analysis showed differences between genders in a few of the biomechanical parameters. Indeed, male rats had ∼33% greater work to fracture than female rats at age 22 wks (p=0.028). In addition, male rats also had a greater ultimate load (∼17%, p=0.044) and elastic modulus (∼61%; p<0.001) at age 28 wks compared to age-matched female rats.

We further evaluated linear relationships between pQCT and biomechanical parameters data. We found several significant positive bivariate correlations in female rats (**Figure 3**) and male rats (**Figure 4**). In female rats, we observed a high positive correlation between cortical area and stiffness (R=0.80), cortical area and ultimate load (R=0.74), cortical thickness and stiffness (R=0.75), cortical thickness and ultimate load (R=0.85), cortical BMD, and stiffness (R=0.76), and cortical BMD and ultimate load (R=0.78) (**Figure 3**).

**Figure 3.**
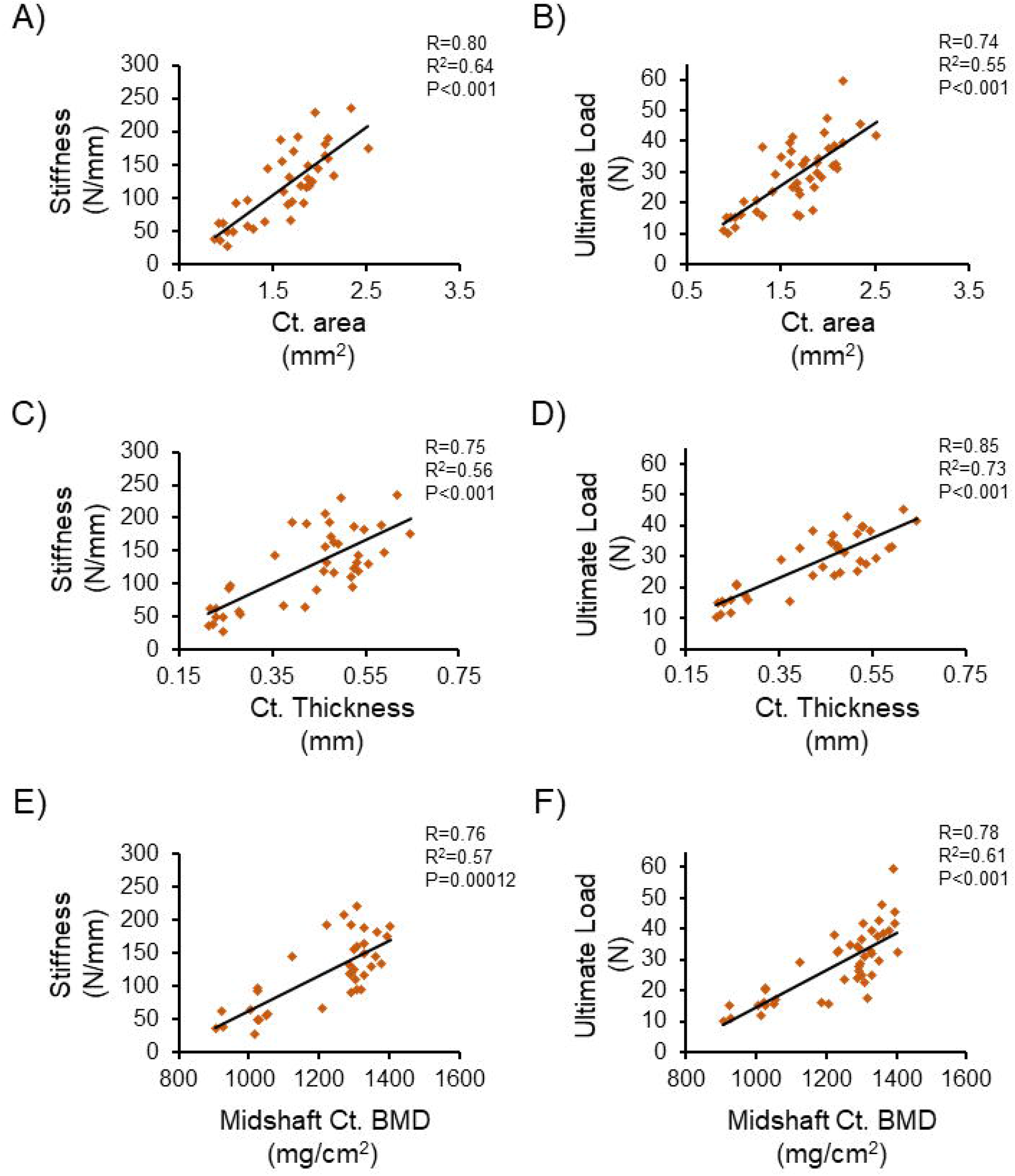
Correlations between biomechanical strength and pQCT parameters at the femoral midshaft of female rice rats. **A**. Stiffness vs. cortical area. **B**. Ultimate load vs. cortical area. **C**. Stiffness vs. cortical thickness. **D**. Ultimate load vs. cortical thickness. **E**. Stiffness vs. cortical BMD. **F**. Ultimate load vs. cortical BMD. Orange diamond markers represent individual female rice rats.

**Figure 4.**
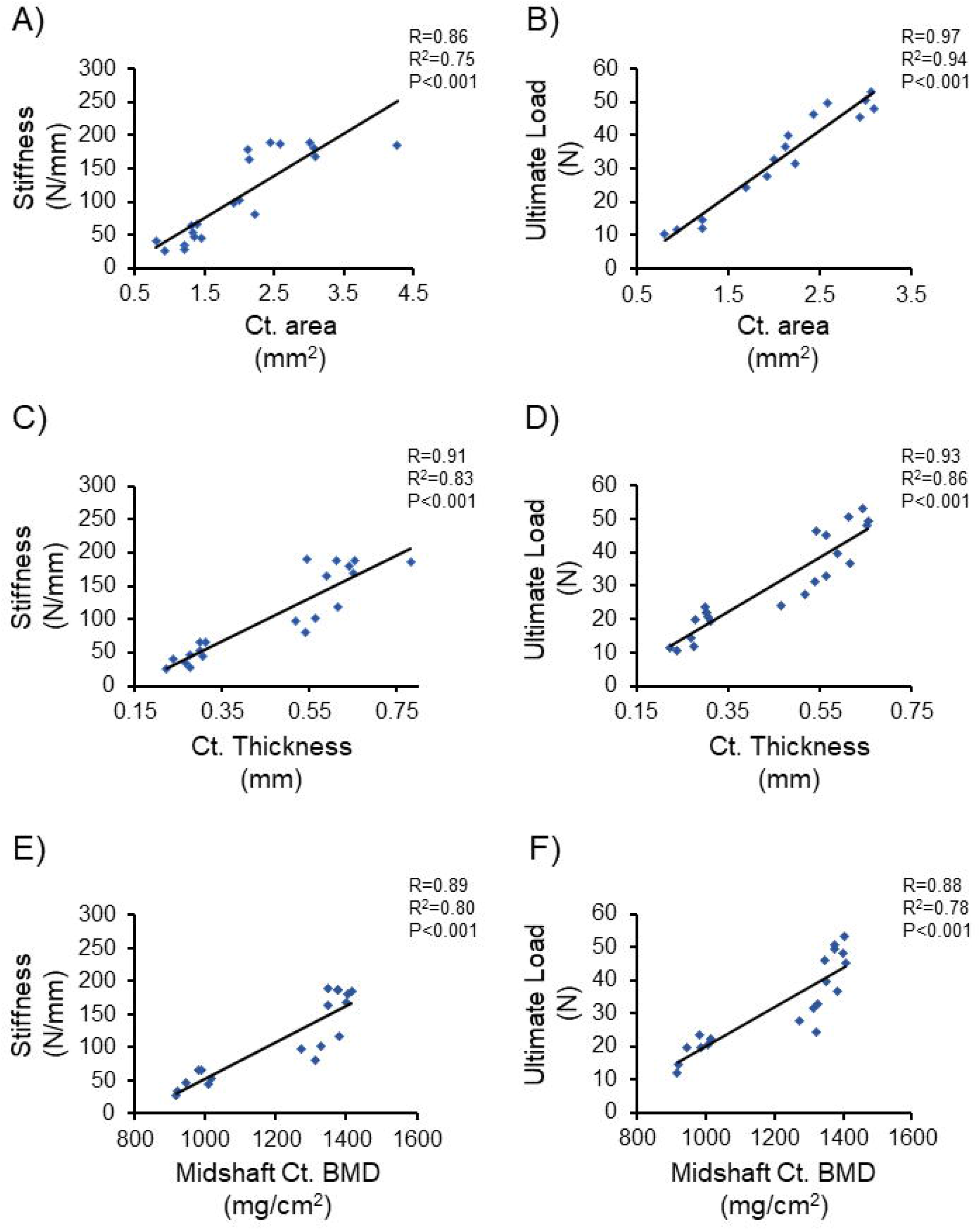
Correlations between biomechanical strength and pQCT parameters at the femoral midshaft of male rice rats. **A**. Stiffness vs. cortical area. **B**. Ultimate load vs. cortical area. **C**. Stiffness vs. cortical thickness. **D**. Ultimate load vs. cortical thickness. **E**. Stiffness vs. cortical BMD. **F**. Ultimate load vs. cortical BMD. Blue diamond markers represent individual male rice rats.

We found a similar trend for these correlations in male rats. In addition, high positive correlations were found between cortical area and stiffness (R=0.86), cortical BMD and stiffness (R=0.89), and cortical BMD and ultimate load (R=0.88). In contrast, positive correlations were found to be very high (0.9-1) between cortical area and ultimate load (R=0.97), cortical thickness and stiffness (R=0.91), and cortical thickness and ultimate load (R=0.93) (**Figure 4**).

### Bone histomorphometry at the proximal tibia metaphysis

At age 4 wks, male rice rats had a greater LGR (∼2-fold) than female rice rats (p<0.05) (**Tables 3 and 4)**. Tibial bone growth dropped abruptly in both genders, reaching similar values at age 10 wks. However, this drop in bone growth between age 4-10 wks was quantitatively more pronounced in male rats (∼6-fold) than in female rats (∼4-fold) (**Tables 3 and 4**). Beyond age 10 wks, LGR values were very low up to age 22-28 wks in both genders.

**Table 3.**
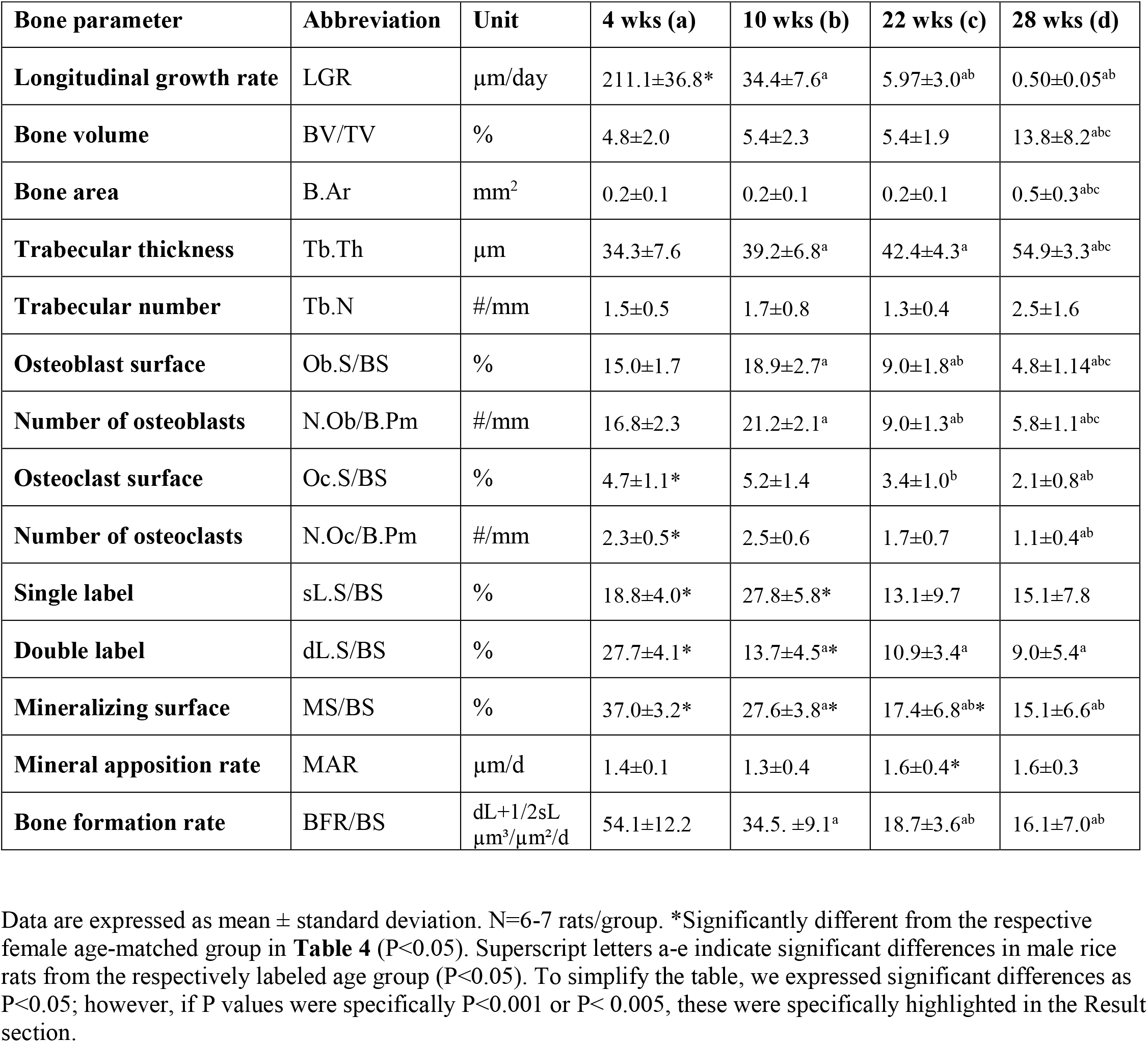
Bone histomorphometry data of the proximal tibial metaphysis in male rice rats distributed by age.

| Bone parameter | Abbreviation | Unit | 4 wks (a) | 10 wks (b) | 22 wks (c) | 28 wks (d) |
| --- | --- | --- | --- | --- | --- | --- |
| Longitudinal growth rate | LGR | µm/day | 211.1±36.8* | 34.4±7.6<sup>a</sup> | 5.97±3.0<sup>ab</sup> | 0.50±0.05<sup>ab</sup> |
| Bone volume | BV/TV | % | 4.8±2.0 | 5.4±2.3 | 5.4±1.9 | 13.8±8.2<sup>abc</sup> |
| Bone area | B.Ar | mm<sup>2</sup> | 0.2±0.1 | 0.2±0.1 | 0.2±0.1 | 0.5±0.3<sup>abc</sup> |
| Trabecular thickness | Tb.Th | µm | 34.3±7.6 | 39.2±6.8<sup>a</sup> | 42.4±4.3<sup>a</sup> | 54.9±3.3<sup>abc</sup> |
| Trabecular number | Tb.N | #/mm | 1.5±0.5 | 1.7±0.8 | 1.3±0.4 | 2.5±1.6 |
| Osteoblast surface | Ob.S/BS | % | 15.0±1.7 | 18.9±2.7<sup>a</sup> | 9.0±1.8<sup>ab</sup> | 4.8±1.14<sup>abc</sup> |
| Number of osteoblasts | N.Ob/B.Pm | #/mm | 16.8±2.3 | 21.2±2.1<sup>a</sup> | 9.0±1.3<sup>ab</sup> | 5.8±1.1<sup>abc</sup> |
| Osteoclast surface | Oc.S/BS | % | 4.7±1.1* | 5.2±1.4 | 3.4±1.0<sup>b</sup> | 2.1±0.8<sup>ab</sup> |
| Number of osteoclasts | N.Oc/B.Pm | #/mm | 2.3±0.5* | 2.5±0.6 | 1.7±0.7 | 1.1±0.4<sup>ab</sup> |
| Single label | sL.S/BS | % | 18.8±4.0* | 27.8±5.8* | 13.1±9.7 | 15.1±7.8 |
| Double label | dL.S/BS | % | 27.7±4.1* | 13.7±4.5<sup>a*</sup> | 10.9±3.4<sup>a</sup> | 9.0±5.4<sup>a</sup> |
| Mineralizing surface | MS/BS | % | 37.0±3.2* | 27.6±3.8<sup>a*</sup> | 17.4±6.8<sup>ab*</sup> | 15.1±6.6<sup>ab</sup> |
| Mineral apposition rate | MAR | µm/d | 1.4±0.1 | 1.3±0.4 | 1.6±0.4* | 1.6±0.3 |
| Bone formation rate | BFR/BS | dL+1/2sL<br>µm<sup>3</sup>/µm<sup>2</sup>/d | 54.1±12.2 | 34.5. ±9.1<sup>a</sup> | 18.7±3.6<sup>ab</sup> | 16.1±7.0<sup>ab</sup> |

Data are expressed as mean ± standard deviation. N=6-7 rats/group. \*Significantly different from the respective female age-matched group in **Table 4** (P<0.05). Superscript letters a-e indicate significant differences in male rice rats from the respectively labeled age group (P<0.05). To simplify the table, we expressed significant differences as P<0.05; however, if P values were specifically P<0.001 or P< 0.005, these were specifically highlighted in the Result section.

**Table 4.** Bone histomorphometry data of the proximal tibial metaphysis in female rice rats distributed by age.

| Bone parameter | Abbreviation | Unit | 4 wks (a) | 10 wks (b) | 22 wks (c) |
| --- | --- | --- | --- | --- | --- |
| Longitudinal growth rate | LGR | $\mu\text{m/day}$ | 94.8±27.8* | 24.8±3.5<sup>a</sup> | 2.7±1.0<sup>a</sup> |
| Bone volume | BV/TV | % | 3.3±1.0 | 9.8.1±7.4<sup>a</sup> | 8.6±3.2<sup>a</sup> |
| Bone area | B.Ar | $\text{mm}^2$ | 0.1±0.0 | 0.2±0.1 | 0.2±0.1<sup>a</sup> |
| Trabecular thickness | Tb.Th | $\mu\text{m}$ | 26.7±4.1 | 35.3±6.8<sup>a</sup> | 54.7±4.3<sup>ab</sup> |
| Trabecular number | Tb.N | $\#/\text{mm}$ | 1.2±0.3 | 2.6±1.7 | 1.6±0.1 |
| Osteoblast surface | Ob.S/BS | % | 13.9±3.5 | 18.1±2.7<sup>a</sup> | 8.0±0.7<sup>ab</sup> |
| Number of osteoblasts | N.Ob/B.Pm | $\#/\text{mm}$ | 18.8±1.5 | 19.5±1.5 | 8.4±1.0<sup>ab</sup> |
| Osteoclast surface | Oc.S/BS | % | 8.1±3.0 | 5.2±1.4<sup>a</sup> | 4.4±1.3<sup>a</sup> |
| Number of osteoclasts | N.Oc/B.Pm | $\#/\text{mm}$ | 3.8±1.4 | 2.7±0.4 | 2.2±0.8<sup>a</sup> |
| Single label | sL.S/BS | % | 29.8±3.6 | 13.5±7.1<sup>a</sup> | 7.8±4.1<sup>a</sup> |
| Double label | dL.S/BS | % | 13.0±4.3 | 7.2±7.4* | 6.3±1.8 |
| Mineralizing surface | MS/BS | % | 27.9±5.8 | 14.0±9.0 | 8.9±3.9<sup>ab</sup> |
| Mineral apposition rate | MAR | $\mu\text{m/d}$ | 1.8±0.4 | 2.3±0.2 | 1.9±0.5 |
| Bone formation rate | BFR/BS | $\frac{\text{dL}+1/2\text{sL}}{\mu\text{m}^3/\mu\text{m}^2/\text{d}}$ | 55.0±15.4 | 38.2±19.5 | 19.1±12.3<sup>a</sup> |

Data are expressed as mean ± standard deviation. N=6-7 rats/group. Superscript letters a-e indicate significant differences in female rice rats from the respectively labeled age group ( $P<0.05$ ). To simplify the table, we expressed significant differences as $P<0.05$ ; however, if P values were specifically $P<0.001$ or $P<0.005$ , these were specifically highlighted in the Result section.

The cancellous bone structural and static histomorphometry data are presented in **Table 3** and **Table 4** for male and female rice rats. Briefly, male rice rats had no differences in BV/TV and bone area among age-groups ≤22 wks. Furthermore, at age 28 wks, these parameters were greater than in rats at age groups ≤22 wks, respectively (p<0.05) (**Table 3). Figures 5 A-D** depict representative photomicrographs of the proximal tibial metaphysis of male rice rats age 4 wks and 28 wks with increased bone mass. Furthermore, trabecular bone thickness progressively increased with age in male rats (p<0.001). However, no significant differences in trabecular number were observed among the different age groups (**Tables 3)**. Osteoblast surface and the number of osteoblasts increased from age 4 wks to age 10 wks (p<0.001). However, after this time point, they progressively decreased with age (**Table 3)**. Osteoclast surface and the number of osteoclasts also tended to be lower in male rats of age-groups ≥22 wks than male rat groups age ≤10 wks (**Table 3**).

**Figure 5.**
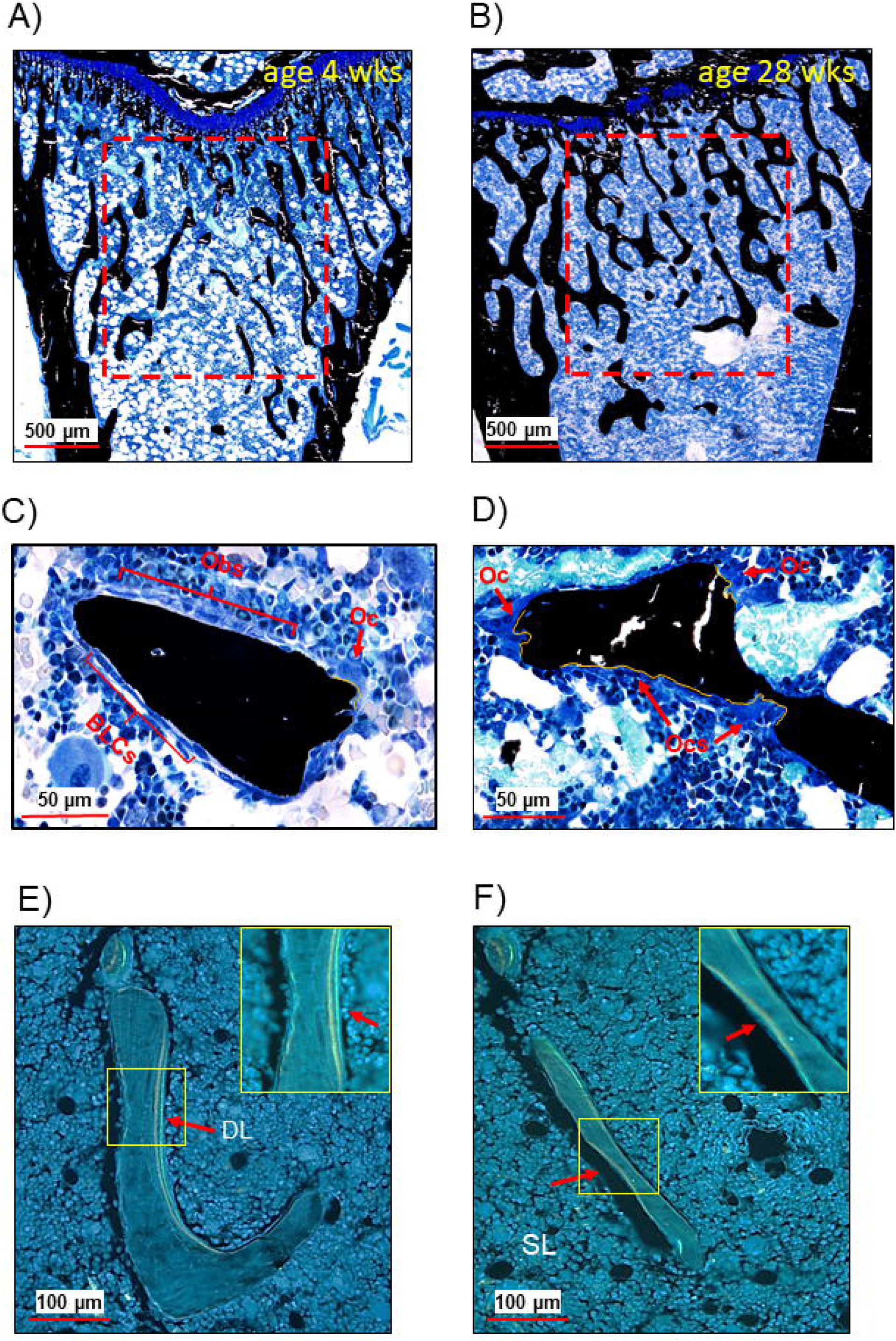
Representative photomicrographs of cancellous bone tissue in the proximal tibial metaphysis of rice rats. **A)** Tibial metaphysis of a male rice rat age 4 wks. (**B**). Tibial metaphysis of a male rice rat age 28 wks. Note the greater amount of cancellous bone (black-stained structures) in the older rat (Von Kossa/tetrachrome stain). **C**. Bone trabecula of a male rice rat age 4 wks depicting numerous osteoblasts (Obs) and bone lining cells (BLCs) on the bone surfaces (red brackets), and a multinucleated osteoclast (Oc) over a small eroded bone surface (yellow irregular line). **D**. Bone trabecula of a male rice rat age 28 wks depicting few osteoclasts (OCs), more extensive eroded bone surfaces (yellow irregular line), and absence of bone-forming surfaces. **E-F**. Dynamic histomorphometry. Fluorescent declomycin (yellow) and calcein (green) labels on bone surfaces. The yellow rectangles on the upper right depict magnified views (4-fold images) of those located in trabeculae. DL: double labels, SL: single label.

In female rice rats, we found greater tibial BV/TV in age-groups ≥10 wks than female rats age 4 wks (p<0.05) (**Table 4)**. As observed with males, female rice rats showed a progressive increase with age in trabecular thickness (p<0.05) but no differences in trabecular number (**Tables 4)**. Furthermore, similar trends as observed in male rats were found in osteoblast and osteoclast parameters in bones from female rice rats (**Table 4**). Indeed, significant reductions in osteoblast surface (p<0.001), osteoclast surface (p=0.009), osteoblast number (p<0.001), and osteoclast number (p=0.007) were observed in female rats age 22 wks compared to female rats age 4 weeks.

Dynamic histomorphometry parameters also tended to decrease with age in male rats (**Table 3**) and female rats (**Table 4**). In male rats, MS/BS was ∼2.5-fold greater at age 4 wks than at age 22 and 28 wks, respectively (P<0.001). Likewise, BFR was ∼3.5-fold greater at age 4 wks than at age 22 wks and age 28 wks, respectively (p<0.001). In female rats, MS/BS (p<0.001) and BFR (p=0.002) were ∼3-fold greater in rats age 4 wks than rats age 22 wks.

**Figures 5 E-F** depict representative photomicrographs of the dynamic histomorphometry evaluation.

## Discussion

This study assessed the rice rat appendicular skeleton’s age- and gender-related features, evaluating quantitative skeletal endpoints using bone histomorphometry, pQCT, and biomechanical testing.

We observed that the body weight and length of femurs and tibiae in rice rats tended to increase with age. However, whereas these parameters continued to rise even at middle age (age 28 wks) in male rice rats, they plateau at age ∼16 wks in female rice rats (early adulthood).

The histomorphometry analysis showed a high LGR in young rice rats (age 4 wks). The high LGR at this young age was particularly remarkable in male rats, with two-fold greater values than in female rats. However, the LGR abruptly dropped from age 4 to 10 wks in both genders and continued falling slower to reach shallow values at age 22-28 wks. Thus, our study shows the growth rate pattern of this species for the first time and suggests that age ∼16 wks is when sexual dimorphism becomes phenotypically apparent, with males’ body size and weight beginning to prevail over females rice rats. Consistent with our findings, male BDIX/Han rats showed a rapid decrease in LGR, nearly zero by age 34 wks (Stark et al., 1996). Other studies also showed a rapid decline in LGR in SD rats, from age 1 to 7 months, followed by a slower rate decline, reaching shallow values by age 12-14 months (Li et al., 1991; Lu et al., 2015). Though a remarkable decrease in LGR, it is well-accepted that the skeleton of rodents continues to grow in length during most of the animal’s life, with differences among genders. In fact, epiphysial growth plates in male rats remain opened beyond age 30 months, whereas in female rats, they tended to close at age 15 months (Jee and Yao, 2001). Interestingly, concurrently with the drastic reduction in LGR that characterizes age progression, Fisher rats show a gradual transition from modeling to remodeling in both cancellous and cortical bone compartments(Erben, 1996).

Femoral total metaphyseal BMC and BMD also increased with age in both genders. However, male rice rats had greater total metaphyseal BMC at age points ≥10 wks and greater total metaphyseal BMD only at age 28 wks than female rice rats. Volumetric BMD is calculated as BMC divided by bone volume. Thus, simply by this mathematical definition alone, and considering the steady increase in BMC observed in both genders with age, the data suggest that female rice rats gain proportionally less metaphyseal bone than male rice rats between age 10 wks and age 22 wks. Likewise, midshaft cortical BMC and BMD, cortical area, and cortical thickness increased with age in both genders. But, again, while these parameters tended to increase steadily to age 28 wks in male rice rats, in the female rice rats, they tended to plateau by age 16-22 wks. Thus, male rice rats have significantly greater values for these parameters than female rice rats at every age point ≥10 wks. As observed in the metaphyses, we found a smaller difference for cortical BMD (P ≤0.05) than BMC (P ≤0.001) among genders at every age point ≥10 weeks. This might be because female rice rats had a proportionally lower cortical bone volume (denominator) than male rice rats.

It has been proposed that BMC, but not BMD, is the more appropriate measure when studying growing animals(Lehtonen-Veromaa et al., 2002; Heaney, 2003). In fact, BMD should never be used in a growth study since there is no reason for true density to change appreciably with growth, and this event rarely happens(Matkovic et al., 1994; Heaney, 2003).

Peak bone mass (PBM) is defined as the amount of bone tissue present at the end of skeletal maturation(Bonjour et al., 1994). It occurs when age-related bone changes are no longer positive, with no further gain in bone mass on average. The PBM coincides with when the skeleton reaches the maximum strength and density(Berger et al., 2010). Though we only measured BMC and BMD in femurs, our study suggests that female rice rats reach PBM at age ∼22 wks (∼5 months). In contrast, our study could not define the age when male rice rats reached PBM because femoral BMC and BMD continued to show a rising pattern by age 28 wks, the latest age point investigated in this study. Studies in rice rats covering a longer age range will be required to define the age at which male rice rats reach PBM and corroborate the female rat findings. Concerning other rodent species, male SD rats achieved PBM at age ∼9 months (Wang et al., 2001). Another study (Iida and Fukuda, 2002) found that cortical and cancellous BMD increased in male Mishima rats between 6 and 12 months of age. However, as in our study, the investigators could not determine PBM because bone density did not stop rising in the rats during the investigation period. Others found that bone mass in female SW-1 mice peaked around midlife (age 12 months) (Bar-Shira-Maymon et al., 1989), and in male C57BL/6J mice, skeletal maturity was reached between age 12-42 weeks (Ferguson et al., 2003). Importantly, significant variations in bone mass have been described among mouse strains(Kaye and Kusy, 1995), indicating that bone mass measures are not only species-specific but also strain-dependent.

Our findings in rice rats also show somewhat equivalent age- and gender-related patterns as previously described in humans (Boot et al., 1997; Gilsanz et al., 2011; McCormack et al., 2017). Indeed, men tend to reach peak bone growth at later time points than females(Alswat, 2017). Furthermore, gender-related skeletal differences become apparent at late puberty in humans, when males, despite comparable body size, show greater BMC and BMD than females (Nieves et al., 2005).

We also observed increased bone stiffness, work to fracture, and ultimate load at the femoral midshaft in older male and female rice rats (age 22-28 wks) compared to younger animals (age 4-10 wks). Thus, our data confirm that independent of gender, the appendicular skeleton becomes stronger and stiffer with age. Furthermore, our study found strong positive correlations between stiffness and ultimate load with the cortical area, cortical thickness, and cortical BMD, suggesting that mechanical properties are dependent on bone quality and bone structure. Age improvements in bone mechanical properties are associated with increased mineralization, bone size, and maturation of the collagen network organization (Wang et al., 2002). In addition, during growth, more mature lamellar bone structures are developed in cortical bone, enhancing bone strength and stiffness (Ferguson et al., 2003).

Overall, our femoral pQCT and strength test findings coincide with those previously reported in other growing rodent species (Danielsen et al., 1993; Iida and Fukuda, 2002). Indeed, female Wistar rats display a rapid increase in femoral length, cortical bone mass, cortical BMC and BMD, and mechanical compressive strength during the first six months of age (Danielsen et al., 1993). Beyond this age, however, these parameters progressed in a more attenuated manner up to age 24 months. Another study looking at age-related changes in bone structure and strength of female and male mice showed a rapid increase in femoral length, cortical bone area, and cortical thickness up to 7 months of age (Willinghamm et al., 2010). The authors also observed increased stiffness and strength with age consistent with changes in bone size and noted significant age-sex interactions for most diaphyseal measurements.

The histomorphometry study found that independent of the gender, young rice rats (age 4-10 wks) have a higher cancellous bone turnover, with increased bone formation and resorption than adult rats (age 22-28 wks). In contrast, mature male and female rice rats showed greater cancellous bone volume and trabecular thickness than young rice rats. This data suggests that despite a significant, progressive decrease in bone cells activity with age and a continuous decay in cancellous bone remodeling, bone formation prevails over bone resorption, reaching a positive bone balance at the bone metaphysis. Similar findings were observed at the metaphysis of growing young Wistar rats, with a turnover rate five-fold higher than in older rats (Baron et al., 1984). Furthermore, greater cancellous bone formation and bone remodeling were found in male SD rats aged 1-3 months but declining at 6 months of age (Lu et al., 2015).

Few studies investigated the hamster’s skeleton’s phenotypic features, a species that belongs to the same family as the rice rat (*Cricetidae*)(Chen et al., 2008). The tibial and femoral cancellous bone volume and cortical bone thickness increased up to age 6 months in this species. However, they significantly decreased from age 12 to 24 months.

Female rice rats are very sensitive to photoperiod(Edmonds et al., 2005). Indeed, when female animals of this species are exposed to short photoperiods, they experienced ovarian function suppression. Furthermore, a long photoperiod is a potent environmental stimulus to increase bone mineral apposition rate in the skeleton of a closed-related species, the Siberian hamsters(Kokolski et al., 2017). Postmenopausal osteoporosis is a condition characterized by decreased bone mass associated with reduced serum levels of estrogens. Taken together, this data suggests that female rice rats that are exposed to reduced light cycles under laboratory conditions would develop clinicopathologic features similar to postmenopausal osteoporosis in women. Thus, the current investigation represents a foundational study to pursue future studies to establish a new model for postmenopausal osteoporosis in a rodent species.

There were a few limitations in this study. One limitation of this study is that we missed the male age 10-16 wks cohort for the strength test analysis and the female age 28 wks cohort for the histomorphometry analysis. Lacking these groups limited gender-specific comparisons for these parameters. Another limitation of our study is that we did not study rats older than age 28 wks old. This limitation prevented us from determining the age for PBM in male rice rats and establishing the age when rice rats of both genders start to show age-related bone loss. A subsequent manuscript will address these critical biologic skeletal stages.

In conclusion, our study in rice rats demonstrates that 1) sexual dimorphism in this species begins to grossly manifest at age ∼16 wks, with significant differences in body size and length of the appendicular bones; 2) cancellous bone growth, bone mineralization, and bone formation rate were significantly greater in younger rice rats but abruptly declined with age; and 3)whereas bone mass, structural cortical parameters, and biomechanical strength rise up to age 16-22 wks in female rice rats, these parameters continued to rise in male rats, at least, up to age 28 wks with significant gender differences at most time points.

## Abbreviations

BMC: bone mineral content
BMD: bone mineral density
LGR: longitudinal growth rate
BV/TV: bone volume
Tb.Th: trabecular thickness
Ob.S/BS: osteoblast surface percent
OCs.S/BS: osteoclast surface percent
MS/BS: mineralizing surface
BFR/BS: bone formation rate

